# Identification of TREK channels in mouse intracardiac ganglion neurons

**DOI:** 10.64898/2025.12.02.691846

**Authors:** A Campos-Ríos, S Herrera-Pérez, M Rodríguez-Castañeda, D Blanco-Remesar, A Rodríguez-Tébar, E De Miguel, A Covelo, JA Lamas

**Author notes:** Corresponding authors: Covelo, A., Lamas, JA. Co-first authors (A Campos-Ríos, S Herrera-Pérez). Co-last authors (A Covelo, JA Lamas).

## Abstract

The intracardiac nervous system is a complex neural network formed by a diffuse plexus, the so-called intracardiac ganglion (ICG). These ganglia locally innervate and control heart function, but little is known about how ICG neuronal activity is regulated. While various ionic currents have been characterized in cardiac tissue, the expression and role of TREK channels in ICG neurons remain unexplored. TWIK-related K^+^ (TREK) channels, members of the two-pore domain potassium channel family, are essential regulators of neuronal membrane resting potential and excitability, with recognized neuroprotective functions. In this study, we have characterized different ionic currents present in mouse ICG neurons, including tetrodotoxin-sensitive Na⁺ currents, delayed K^+^ rectifiers, Ca^+2^-dependent K⁺ currents and A, H and M currents. Notably, we provide the first evidence of functional TREK channel expression in these neurons. Our findings suggest that TREK channels contribute to excitability control and cellular homeostasis in ICG neurons, underscoring their potential as therapeutic targets for cardiac disorders.

**KEY POINTS:**

- This study provides the first evidence of functional TREK channel expression in ICG neurons.
- Comprehensive profiling of ionic currents shaping excitability in ICG neurons.
- Neuroprotective TREK channels might emerge as potential therapeutic targets for cardiac disorders.

## INTRODUCTION

The intracardiac nervous system (ICNS) is a diffuse ganglionated plexus formed by a network of ganglia (known as the intracardiac ganglia, ICG) and interconnecting nerves^1^. The ICG form a complex structure comprising sensory, motor, and local regulatory neurons that express multiple neurochemical phenotypes^2^ and are the central hub of the integrative autonomic network regulating heart function^3^. Located in the adipose tissue of the atria and in the atrioventricular (AV) groove, it plays a crucial role in the regulation of cardiac conduction velocity, inotropism and heart rate^1,2,4–7^.

In the past years, several studies have characterized the electrophysiological properties of the mammalian ICG neurons using guinea pigs and rats^8–13^. These studies have depicted some ion channel underlying the ICG neuronal excitability and action potential (AP) shaping, but whether these ion channels are expressed in other species is currently unknown. Additionally, no one until date have studied the expression of TWIK-related K^+^ (TREK) channels in the intracardiac nervous system. These channels, composed by TREK1, TREK2 and TRAAK, are members of the two-pore domain potassium (K2P) channels family and are widely expressed in the central, peripheral and autonomic nervous systems (CNS, PNS and ANS, respectively)^14–16^. They are key controllers of neuronal resting membrane potential (RMP) and excitability^17^ due to their unique features. They can be modulated by many physical and chemical stimuli, and many pharmacological substances influence their activity^18–21^, suggesting that they may be involved in plenty of physiological processes. In fact, recent studies have implicated cardiac TREK channels in pathologies such as arrhythmia, atrial fibrillation (AF) or long QT syndrome^22–24^. Additionally, TREK channels are activated by antiarrhythmic agents^25–27^, suggesting that they may play a central role in heart function. Several studies have demonstrated the presence of TREK channels in the heart cardiomyocytes and nodal cells of different species, including mice and human^22,28–32^. However, their expression and function in ICG neurons remains unexplored.

Here, we have characterized whether mouse ICG neurons express different ion currents, including tetrodotoxin (TTX)-sensitive Na⁺ currents, delayed rectifiers, Ca^+2^-dependent K⁺ currents, as well as A, H and M currents (I_A_, I_H_ and I_M_, respectively). Furthermore, we provide the first evidence that ICG neurons have TREK-like currents and demonstrate the expression of all three TREK channel members in ICG neurons. These findings suggest that the expression of ion channels in ICG neurons may be conserved among species and that TREK channels contribute to the regulation of neuronal excitability and homeostasis in the intracardiac nervous system.

## MATERIALS AND METHODS

### Experimental subjects

Animals were housed in the Bioexperimentation Centre of the University of Vigo. All experimental procedures were performed using male and female 30 – 90 days CD-1 mice. The animals had access to commercial food and water *ad libitum* and were maintained under controlled temperature (20 – 24 °C) and relative humidity (45 – 55%) with 12h light/dark cycles. Animal protocols were approved by the European (2010/63/EU) and Spanish Research Council (RD 53/2013), as well as by the Scientific Committee of the University of Vigo, following the European and Spanish animal protection directives.

### Heart extraction and sample preparation

Animals were sacrificed in a euthanasia chamber with a gradual increment of CO_2_ and death was confirmed by cervical dislocation. A thoracotomy was performed, and the myocardium was perfused with saline serum 0.9% to eliminate the blood. Then, the whole heart was cut off and placed in a 35-well dish filled with Leibowitz (L-15) medium, where, under a dissection magnifying glass, the dorsal region was isolated. Mouse ICG are mostly located in the central portion of the atrium, so this region was dissected before separating the atria appendages^3^

### Primary neuronal cultures

After the heart dorsal region was isolated and minced under sterilized conditions, the sample was incubated in a collagenase solution (3mg/ml, 35 min at 37 °C and 5% CO_2_) followed by an incubation in a trypsin solution (1 mg/ml 35 min at 37 °C and 5% CO_2_). For immunocytochemical techniques, DNase I (0.25 mg/ml) was added to the trypsin solution. After enzymatic disaggregation, the sample was mechanically disaggregated until a single cell suspension was obtained. Next, the suspension was transferred to a conical tube filled with Ficoll-Paque solution and centrifugated (1600 rpm at 22 °C for 15 min) to eliminate erythrocytes and dead cells. Then, the interphase was collected and centrifuged again (1600 rpm for 3 min). The pellet was finally resuspended in culture medium, and neurons were plated into 35 mm dishes previously treated with laminin (10 µg/ml for 2 hours at 37 °C and 5% CO_2_). They were maintained in these conditions during the following 24 hours. The primary cultures used for immunofluorescence were directly plated onto coverslips treated with laminin as described above.

### Nissl staining

The heart dorsal region was fixed in buffered neutral formalin (BNF) for 24 h at 4 °C, dehydrated in a battery of alcohol solutions (H_2_O - 70 ° - 80 ° - 90 ° - 96 ° - 100 °) and included in paraffin for 24h. Then, the tissue was cut in 10 µm sections in a microtome (Leica, Germany) and serial cuts were placed in slides. Afterwards, the sections were deparaffinated, hydrated and stained for 5 minutes with Cresyl violet, staining that shows a clear proclivity for neuronal bodies. The samples where again dehydrated, coversliped and visualized with an ECLIPSE Nikon Ni-E microscope. Nikon NIS-Elements software and ImageJ software were used to process the images.

### Immunohistochemistry and immunocytochemistry

The immunohistochemistry (IHC) technique was performed in both tissue and ICG neuronal primary culture preparations. For IHC in whole tissue, the dorsal heart region was included in optimal cutting temperature compound (OCT, CellPath, United Kingdom) embedding matrix and submerged in liquid N_2_ for its total and fast freezing. Hearts were serial cut into 10 µm slices and placed in gelatinized slides. Slides were placed at −20 °C for 24 hours to promote adhesion. Samples were fixed with acetone 3%, dried for 30 minutes and hydrated. After washing the samples with phosphate buffered saline (PBS), unspecific sites were blocked with bovine serum albumin (BSA) 1% for 1 hour. Then, the slides containing the samples were separately incubated in the following primary polyclonal antibodies: rabbit anti-calretinin, rabbit anti-tyrosine hydroxylase, rabbit anti-KCNK1, anti-KCNK2, anti-KCNK10 (Alomone Labs, Israel), 1:100 diluted in Envision Flex Antibody Dilution (Dako, Denmark) and incubated overnight at 4 °C in a humidity chamber. DAPI (Thermo Fisher Scientific, USA) was utilized to stain neuronal nucleus. Samples were then incubated in Envision Flex Antibody Dilution with secondary polyclonal antibody: goat anti-rabbit AlexaFluor555 (Thermo Fisher Scientific, USA) for 1 hour at room temperature (RT). Finally, slides were dried and covered with a coverslip with Prolong Gold Antifade Mountant (Thermo Fisher Scientific, USA).

For immunocytochemistry, ICG neuronal primary cultures were first fixed with 2% paraformaldehyde (PFA) for 10 minutes at 37 °C and unspecific sites were blocked with 3% BSA for 1 hour at RT in a shaker. Afterwards, samples were incubated with the above cited primary antibodies (1:500 dilution) overnight at 4 °C. Cells were then washed with PBS and incubated with the secondary antibody listed above for 1 hour at RT and the coverslips were mounted in a slide using a drop of Prolong Gold Antifade Mountant (Thermo Fisher Scientific, USA).Samples were visualized with Leica Stellaris 8 100.0 × 3.5 OIL objective controlled by Leica LAS X software (Leica Microsystems, California, USA) at the Galicia Sur Biomedical Foundation facilities and Leica LAS X software and ImageJ software were used to create microscopy figures.

### Electrophysiological recordings

Whole-cell patch-clamp recordings (perforated patch variant) were performed 24 hours after primary cell cultures were established. Artificial cerebrospinal fluid (ACSF) was perfused at approximately 10 ml/min. The ACSF contained (in mM): NaCl 140, KCl 3, MgCl_2_ 1, CaCl_2_ 2, D-glucose 10, HEPES 10, gassed with synthetic air and a pH of 7.2 adjusted with Tris (Tris (hydroxymethyl)-amino-methane) (osmolarity 270-300 mOsm/L). Patch pipettes with resistances between 4 and 7 MΩ were fabricated from borosilicate capillaries (Harvard Apparatus, Massachusetts, USA) using a P-1000 horizontal puller (Sutter Instrument, California, USA). Patch pipettes were backfilled with intracellular solution containing (in mM) KAc 90, KCl 20, MgCl_2_ 3, CaCl_2_ 1, EGTA 3, HEPES 40, NaOH 1 to adjust pH to 7.2 (osmolality 270-300 mOsm/L). Amphotericin-B (75 µg/ml) was included to the intracellular solution for performing the perforated patch variant. Signals were acquired with an Axon amplifier and the pClamp software (Molecular Devices, California, USA). Signals were acquired at 10KHz and filtered at 5 kHz for current-clamp experiments and acquired at 5 kHz and filtered at 1 kHz for voltage-clamp experiments. For single-channel experiments, data was sampled at 20 KHz and low-pass filtered at 2 KHz using the amplifier built-in filter. Clampfit 10 (Molecular Devices, California, USA), OriginPro 8.5 (OriginLab Corporation, Massachusetts, USA) and GraphPad Prism 9 (GraphPad Software Inc., California, USA) softwares were used for data analysis, statistics and representation. For experiments assessing firing rate of ICG neurons, current pulses from −50 to 175 pA (1 second each, 25 pA increment) were applied in current clamp mode. Individual AP properties were measured as follows: latency was measured from the beginning of the stimulus to the AP positive peak; amplitude was determined from the most positive to the most negative voltage of the AP; duration is the 50% amplitude width; after-hyperpolarization (AHP) was calculated from the resting potential to the most negative voltage; and finally, frequency was calculated between the first and the second AP fired.

To study Na_v_ and K_Ca_ currents (I_Nav_ and I_KCa_, respectively), in voltage-clamp configuration jumps from −50 to 10 mV (20 ms) were applied in the absence and the presence of tetrodotoxin (TTX) 0.5 μM and CdCl_2_ 100 μM. Na_v_ and K_Ca_ currents were obtained by subtracting signals obtained in the presence of TTX and CdCl_2_ respectively, from control signals.

For A current (I_A_) and delayed rectifiers (I_KV_), a jump from −120 mV to 10 mV (20 ms) was followed by another jump from −50 to 10 mV (20 ms). In the first jump both currents of study are overlapped, but in the second jump only the I_KV_ is activated, so I_A_ can be subtracted.

To study the presence of I_M_ and I_H_ in ICG neurons, ramps from −30 to −100 mV (10 mV/s rate) were performed in voltage-clamp mode before and after tetraethylammonium chloride (TEA) 15 mM or CsCl 1 mM application to unmask I_M_ and I_H_, respectively.

For experiments assessing the effect of TREK current activators, neuron membrane potential was held at −30 mV in voltage-clamp mode. TREK currents were isolated with a cocktail of ion channel blockers containing TTX 0.5 μM, CdCl_2_ 100 μM, CsCl 1 mM, TEA 15 mM, 4-Aminopyridine (4-AP) 2 mM, apamin 200 nM, paxilline 1 μM and clemizole 10 μM. Conductance was monitored by applying pulses of −15 mV every 50 ms. Single-channel recordings were performed in the cell-attached configuration. Bath and pipette solutions were identical and contained (mM): KCl 150, MgCl_2_ 1, EGTA 5 and HEPES 10, pH 7.2. The electrode resistance ranged from 10 to 12 MΩ. Single-channel openings faster than 50 µs were discarded. Threshold detection for single-channel openings was set at 50 % of the amplitude. Voltage steps from −100 to 100 mV were applied and the currents acquired in every step were used to perform a voltage-intensity curve (I-V) and characterize the channel. By fixing the membrane at −60 mV, amplitudes of each channel were measured using pClamp software (Molecular Devices, California, USA) and conductance was calculated (*V= I * 1/g*, where *g* is conductance). All compounds were purchased in Sigma-Aldrich (Madrid, Spain), except for TTX, from Tocris Bioscience (Abington, United Kingdom).

### Statistics

Results are represented as mean ± standard error of the mean (SEM). The Shapiro-Wilk test was performed to determine normality. ANOVA test was applied to study differences between three firing groups. Paired t-Student test was performed in voltage clamp experiments to compare the difference between conditions to unmask drug-sensitive currents. The differences among groups were considered significant when p<0.05 (*), p<0.01 (**) or p<0.001 (***). Dose-response curves were fitted using the Hill equation: *y=START+(END-START)x^N/(K^N+x^N)*, where *K* corresponds to the EC_50_ and *N* is the Hill coefficient. A Hill coefficient > 1 was considered as indicative of multiple binding sites, while < 1 is indicative of a single receptor-molecule binding site (Monod et al., 1965; Weiss, 1997).

## RESULTS

### TH and CR reactivity in ICG neurons

To study the histological localization of ICG neurons, hearts were embedded in paraffin (n=6), sectioned with a microtome (Figure 1 A) and stained using Nissl technique, which stains neuronal bodies. Mouse ICG are diffuse plexuses, consistent with that, we observed neuronal staining localized in close proximity to the AV groove and the atrial adipose tissue in all the hearts studied (Figure 1 B). The stained ganglia were heterogeneous in shape and size consisting of 15-60 neurons that were oval or round and had prominent somas, and smaller satellite glial cells (Figure 1 C). Large, rounded nuclei with weaker DAPI were observed in neurons compared to those from putative satellite glial cells, whose nuclei were smaller and oval shaped. Based on this distinction, DAPI staining was subsequently used as an approximation to localize neurons within the ganglia. Cells that showed weaker DAPI staining were considered putative neurons, while the strong DAPI staining was identified as satellite glia (Figure 1 E-H).

**Figure 1.**
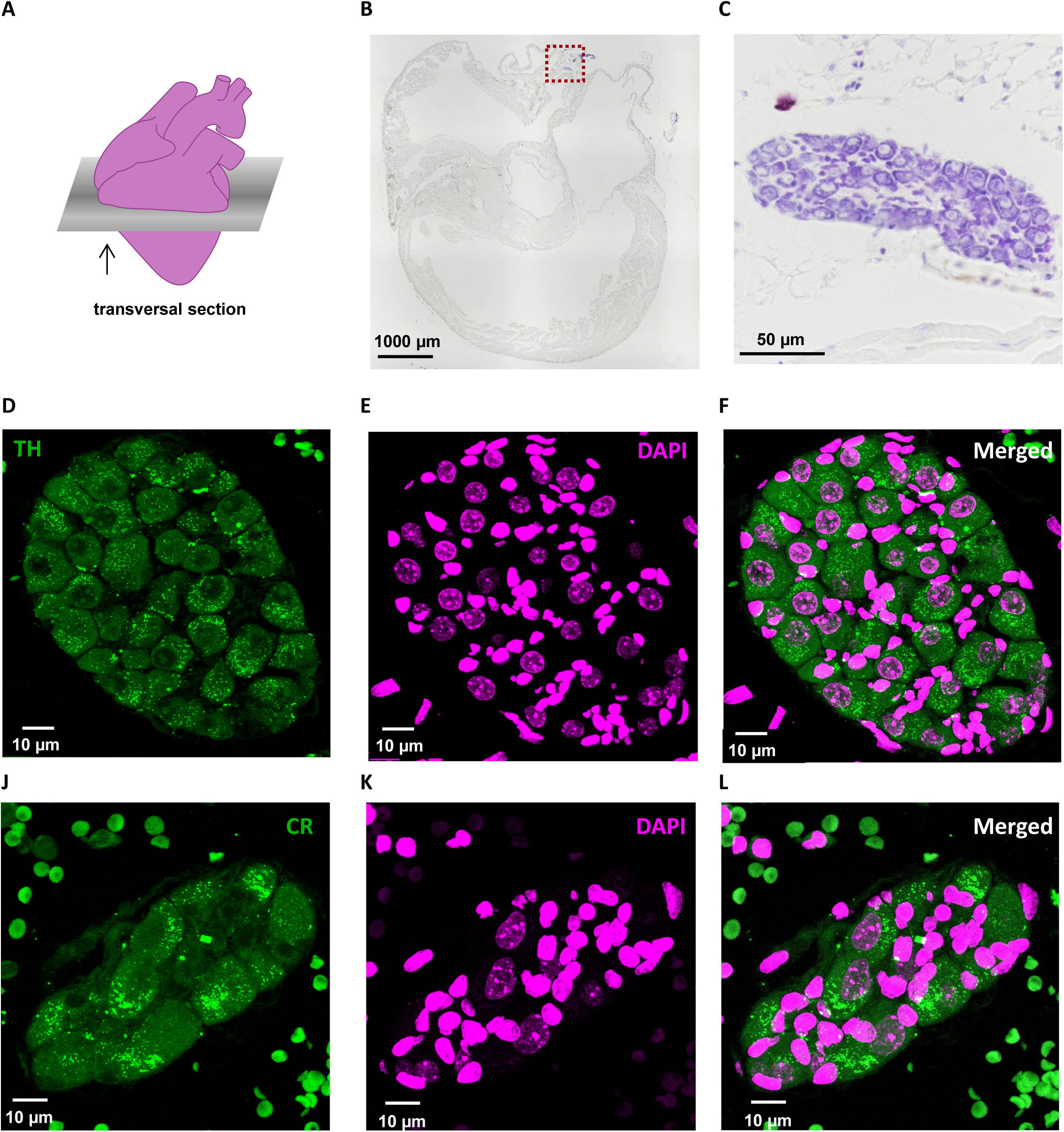
Histological characterization of the mouse intracardiac ganglion. (A) Schematic representation of the cardiac transversal tissue sections. (B) Representative Nissl staining image of a mouse heart section containing the intracardiac ganglion (ICG) located above the atrioventricular groove (square). (C) Magnified imaged of the ICG shown in (B) (40X magnification). (D-F) Tyrosine-hydroxylase (TH) and (G-I) calretinin (CR) positive neurons (green) located in the ICG with nuclei are stained with DAPI (blue).

We next studied the histological properties of ICG neurons. To do this, hearts were embedded in optimal cutting solution and directly preserved in liquid nitrogen (n=3) and cut in a cryostat. After serial sectioning of the whole heart, some of the slides were Nissl stained to localize the ICG neurons and adjacent sections were subsequently used for studying the expression of specific markers using IHC. The ICG receives catecholaminergic innervation^2^ and many studies have reported tyrosine hydroxylase (TH) reactivity in ICG neurons as a basic neurochemical phenotype^3,33^. Consistent with this, we found TH expression in ICG neurons in all the samples checked (n=3, Figure 1 D-F). We further found that calretinin (CR), a calcium-binding protein involved in Ca^+2^ signalling and homeostasis^34^, is expressed in ICG neurons (n=3, Figure 1 G-I). Although this protein is known to be widely expressed in the ANS^35–37^, this is the first time that its expression is demonstrated in the ICNS.

### Electrophysiological characteristics of ICG neurons

Perforated whole-cell electrophysiological recordings were performed to establish the passive and active properties of ICG neurons. In current-clamp mode, a RMP of −70±1.2mV (n=45) (Figure 2 A) and a membrane capacitance of 33.9±1.4 pF (n=45) (Figure 2 B) were observed.

**Figure 2.**
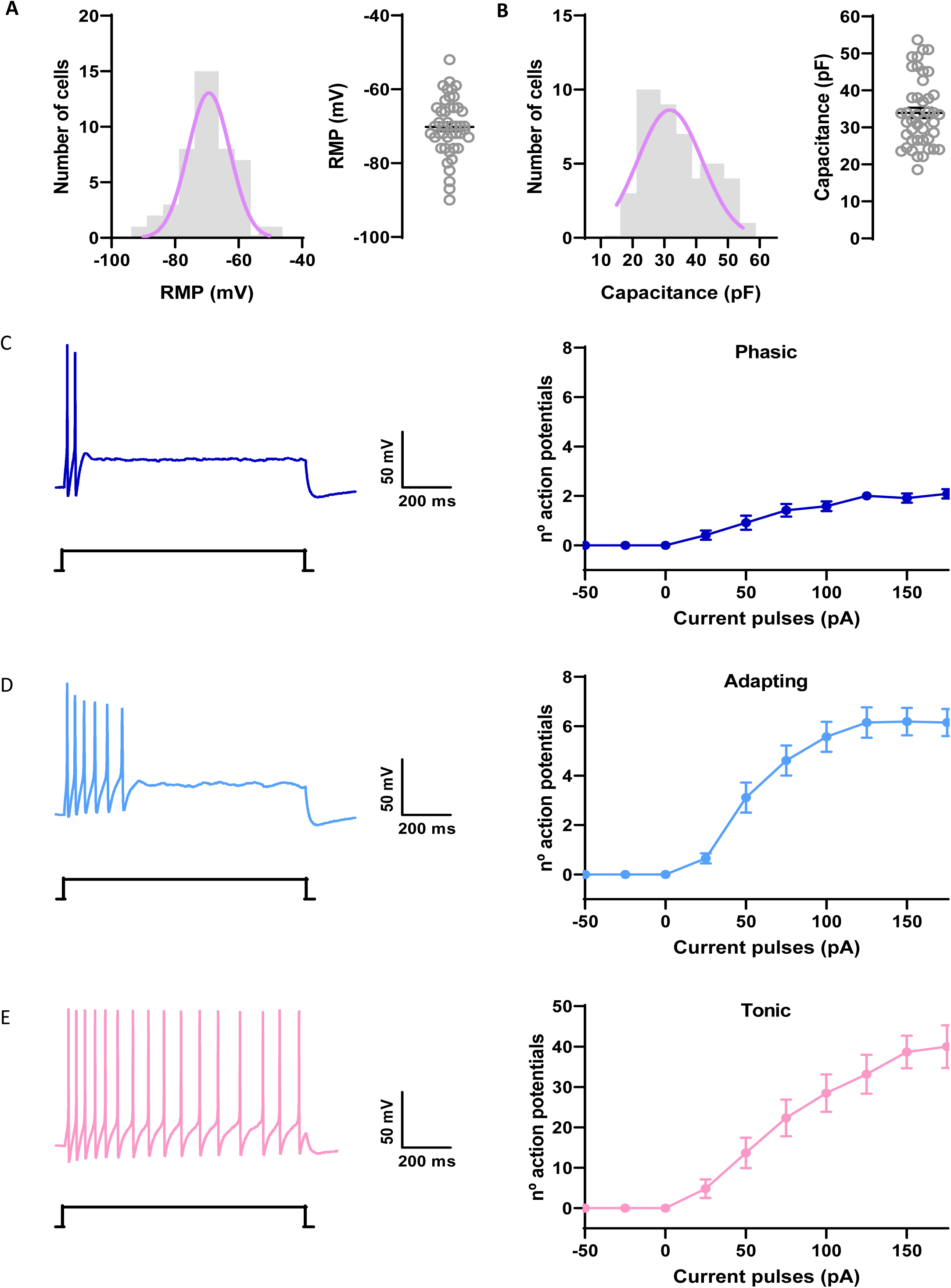
Intracardiac ganglion neurons intrinsic properties. (A) Distribution of the resting membrane potential (RMP; left) and average RMP (right) recorded from ICG neurons. (B) Distribution of the membrane capacitance (left) and average membrane capacitance (right) recorded from ICG neurons. (C) Representative traces from a phasic neuron showing the AP firing in response to a depolarizing step (left) and number of AP triggered by increasing depolarizing steps (right). (D) and (E) like (C) but for adapting and tonic neurons, respectively. Data are expressed as mean ± SEM.

By fixing the membrane potential at −60 mV and applying one second current pulses from −50 to 175 pA (increments of 25 pA), three different firing patterns were observed in IGC neurons: 24% of the neurons (n=12 out of 50 neurons) displayed a phasic adapting discharge of 1 to 3 AP (phasic neurons, Figure 2 C), whereas the 66% (n=33 out of 50 neurons) presented an adapting firing (adapting neurons, Figure 2 D), discharging a maximum AP firing of 5.47±0.6 AP. The remaining 10% (n=5 out of 50 neurons) discharged AP during the whole current pulse (tonic neurons, Figure 2 E), with a maximum AP firing of 42.2±5.6.

AP latency, amplitude, duration, frequency and afterhyperpolarization (AHP) were measured. No differences were found in RMP, capacitance, latency, amplitude, duration and AHP between the three neuronal subgroups (Figure 3). On the contrary, tonic neurons displayed a higher instantaneous frequency (51,4±2,9, n=5) than adapting (29,2±1,4, n=23) and phasic (31,1±5,5, n=10) neurons (Figure 3 H, One-way ANOVA, p<0.001), without differences in AP firing frequency between these two last neuronal types.

**Figure 3.**
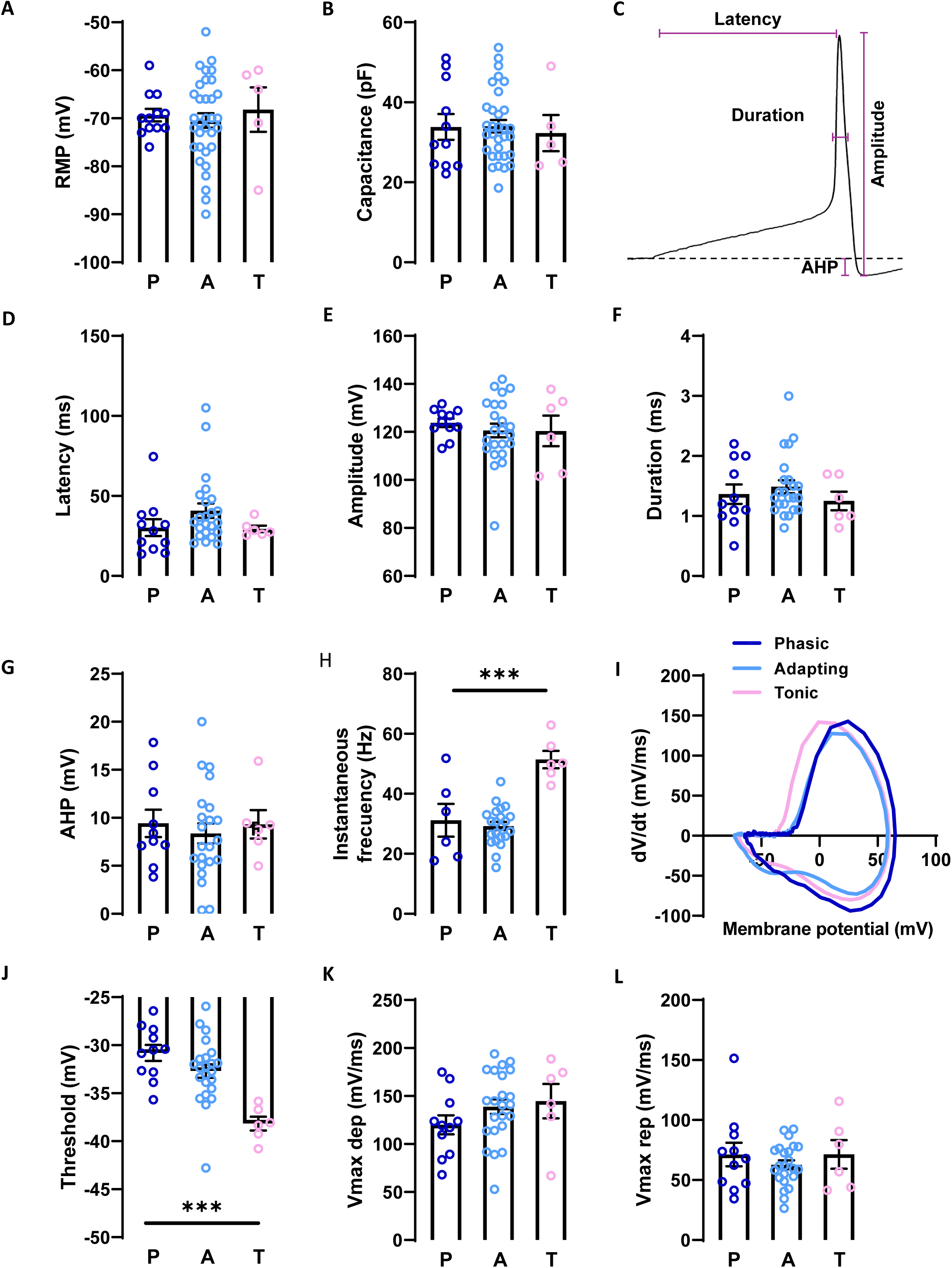
Characteristics of individual action potentials. (A) Resting membrane potential (RMP) and capacitance were similar in phasic, adapting and tonic ICG neurons. (B) As observed in the scheme, (C) AP latency, AP amplitude, AP duration, (D) afterhyperpolarization (AHP) and AP instantaneous frequency were measured in phasic, adapting and tonic ICG neurons. (E) Phase diagram of representative AP recorded in phasic, adapting and tonic ICG neurons. (F) AP firing threshold, maximal velocity of depolarization (V max dep) and a maximal velocity of repolarization (V max rep) measured in the three neuronal types. Data are represented as mean ± SEM; significance is represented as (*) p<0.05, (**) p<0.01, (***) p<0.001. P: phasic; A: adapting; T: tonic.

To assess the AP discharge velocity, the derivative of the voltage with respect to time was plotted against the membrane potential (Figure 3 I). Although no differences were observed in maximal velocity of depolarization and maximal velocity of repolarization, we found that the AP threshold of tonic neurons is significantly lower (−38,2±0,7, n=6) than that of phasic (−30,8±0,8, n=12) and adapting (−32,7±0,7, n=33) neurons (Figure 3 J).

### Sensitivity to TTX, delayed rectifiers, A current and Ca^2+^-dependent potassium currents

We next studied the type of ionic channels expressed by ICG neurons. Voltage-gated Na^+^ channels are blocked by TTX; however, several cell types, including some ANS neurons^38–40^, express TTX-resistant Na^+^ currents. Thus, we explored the presence of these currents in ICG neurons. To do that, a voltage step (from −50 to 10 mV) was applied in the presence and absence of TTX (Figure 4 A). This step induced an inward current (I_Na_: 1420.61 ± 186.01 pA, n=15, Figure 4 A and B) that was fully blocked by TTX, suggesting that ICG neurons do not express voltage-gated Na^+^ channels insensitive to TTX as previously proposed^3^. Additionally, the I_Na_ current was followed by an outward current (I_KCa_: 267.41 ± 71.46 pA, n=15, Figure 4 A and B) that was blocked by CdCl_2_, indicating that these neurons express Ca^2+^-dependent K^+^ channels, as previously demonstrated in mouse, rat and guinea pig ICG neurons^3,11,12^.

**Figure 4.**
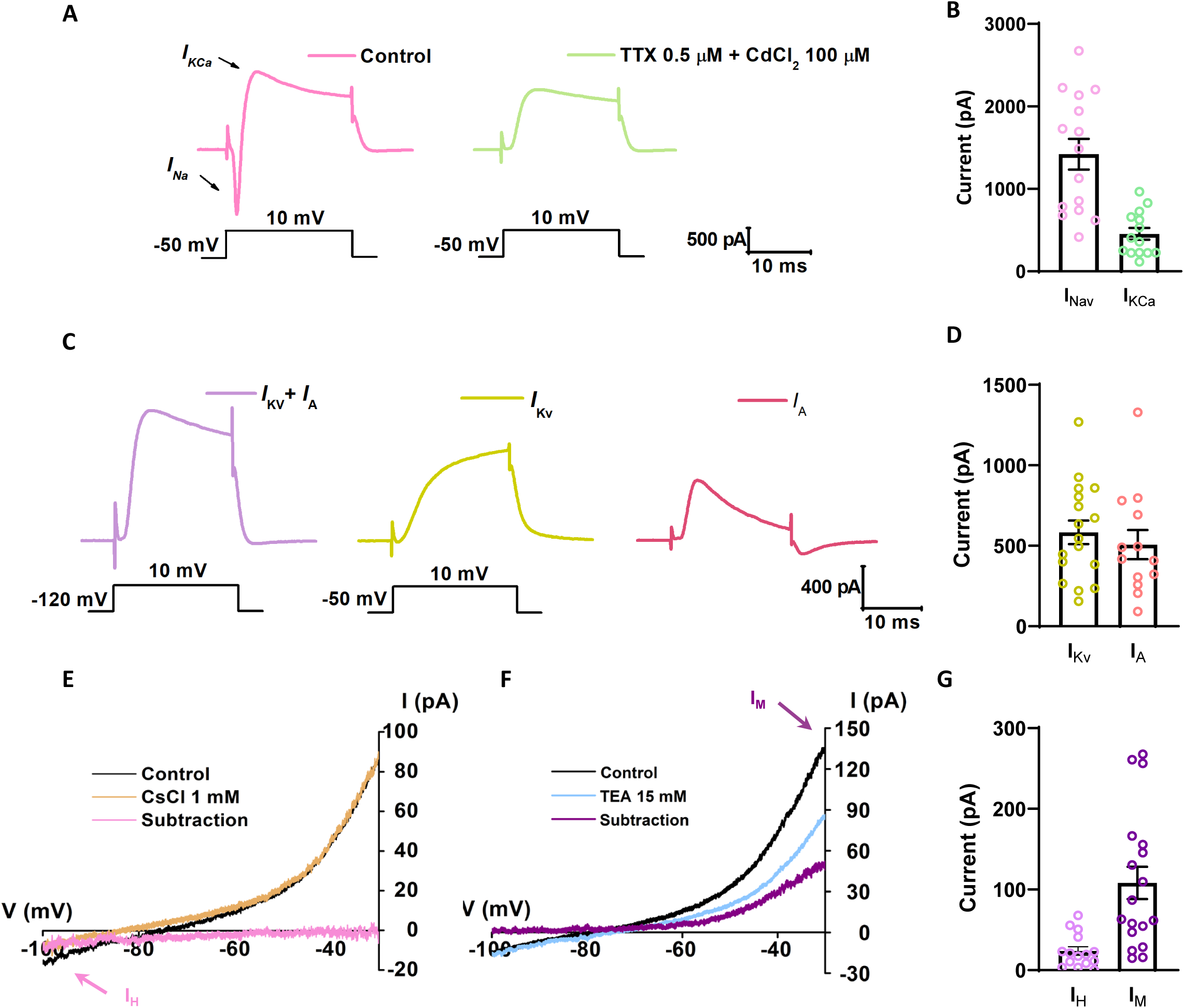
TTX sensitive and different potassium currents participate in ICG neuron electrical activity. (A) Representative traces showing a TTX-sensitive Na^+^ current (I_Na_) and a Ca^+2^-dependent K^+^ current (I_KCa_) recorded in cultured ICG neurons before (pink) and after (green) TTX and CdCl_2_ application. (B) Average amplitude of I_Na_ and I_KCa_. (C) Representative traces of delayed rectifiers (I_KV_) and A-type current (I_A_) together (Ca) when applying a current step from −120 mV to 10 mV. Consecutively, applying a pulse from −50 to 10 mV only activates I_KV_ (Cb), and I_A_ can be observed subtracting I_KV_ to the first jump protocol (Cc). (D) Average amplitude of I_KV_ and I_A._ (E) Representative ramps obtained before and after CsCl 1mM application and its subtraction (pink, H current). (F) Representative ramps obtained before and after TEA 15 mM application and its subtraction (green, M current). (G) Average amplitudes of H current (I_H_) and M current (I_M_). Note: I_H_ was measured in the subtraction trace at −100 mV and I_M_ was measured in the subtraction trace at – 30 mV.

Furthermore, the presence of other voltage-gated currents such as I_KV_ and I_A_ has been in rat and guinea pig ICG neurons^3,41–43^. To assess the presence of these currents in mouse ICG neurons, two consecutive voltage steps were performed. The first step (from −120 to 10 mV) induced an outward current composed by both both I_A_ and I_KV_ currents (Figure 4 C), and the second (from - 50 to 10 mV) activated only I_KV_ currents (583.54 ± 73.2 pA, n = 17, Figure C and D). By subtracting these two currents we could calculate the I_A_ (507.38 ± 90.8 pA, n = 13, Figure 4 C and D). These results show that mouse ICG neurons express potassium A-type channels and K^+^ delayed rectifier channels.

Next, the Cs-sensitive I_H_ current was unmasked using a voltage clamp ramp from −30 to −100 mV (10 mV/s rate) in the presence of TTX 0.5 µM and CaCl_2_ 100 µM. The ramp was performed before and after CsCl application (1mM) and I_H_ was calculated by subtracting the signal obtained in the presence of CsCl to the one obtained in control conditions (Figure 4 E). By doing this, we found that I_H_ begins to open below −70 mV and reached −5.75 ± 2.10 pA at −100 mV (n = 10, Figure 4 E and G), indicating that ICG neurons express hyperpolarization-activated cyclic nucleotide-gated (HCN) channels.

We, then, repeated this protocol before and after TEA application (15mM) to reveal the presence of TEA-sensitive I_M_ current (Figure 4G). We observed I_M_ current in ICG neurons for the first time (Figure 4 F). I_M_ reached 108.19±20.02 mV at −30 mV (n= 18, Figure 4 F and G) and was completely closed below −60 mV^44–46^. These data indicate that ICG neurons express M current, presumably due to the expression of KCNQ2 and 3 channels^47,48^.

### TREK-like currents are present in mouse ICG neurons

Next, we studied whether ICG neurons express TREK channels. For this, neurons were recorded in voltage clamp mode (V_hold_ = −30 mV) and TREK currents were isolated by using a cocktail of blockers of other ionic channels (see methods). Riluzole, a neuroprotective drug indicated for amyotrophic lateral sclerosis treatment^49^ and considered a TREK channel activator^20,21,38^, was applied in the bath. Different riluzole concentrations were added to the bath solution (3 µM, 10 µM, 30 µM, 100 µM, 300 µM and 1000 µM, Figure 5 A) and an outward current was generated. The values of this dose-response curve were adjusted to a Hill equation, where the half maximal effective concentration was EC_50_ = 126.20 ± 42.97 µM with a Hill coefficient > 1. Furthermore, an increase in the conductance (Figure 5 B) from 4.48±0.56 nS to 6.67±0.65 nS was observed with the addition of riluzole 300 µM (n=12, p<0.001).

**Figure 5.**
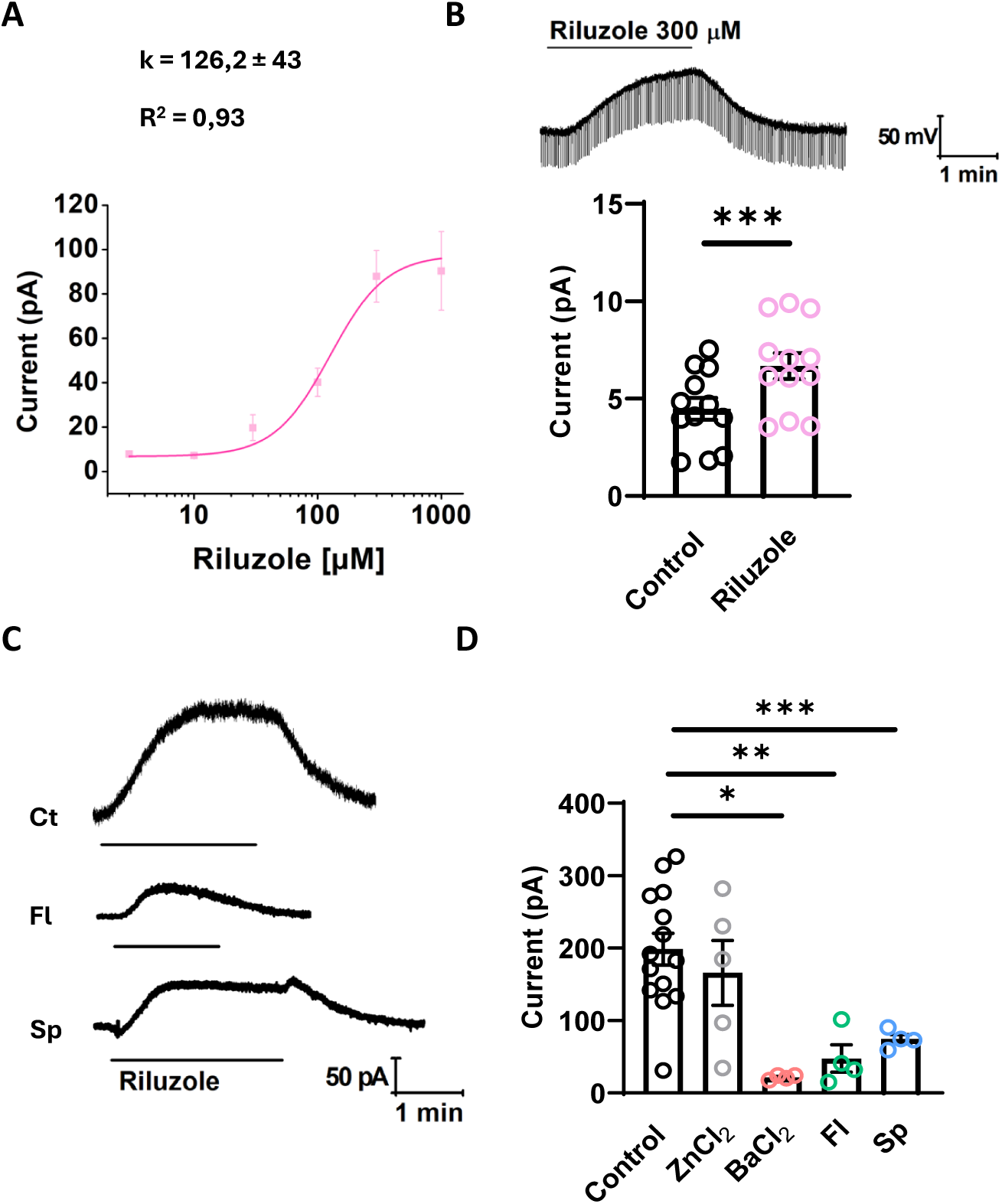
TREK channel blockers reduce the riluzole activated current. (A) Riluzole-activated current is dose-dependent, with a EC50 (k) of 126.2±43 µM (adjusted to Hill equation, R2 = 0,93) and an increase in conductance (B) when riluzole 300 µM is added to the bath solution (paired t-Student, n=12, p<0.001). In the presence of the blockers cocktail to isolate TREK currents, riluzole was applied to the bath (control, C). Riluzole activated current, despite not being reduced by Zncl2, a Ca2+ channel blocker, was significantly diminished by BaCl2, fluoxetine and spadin (D), all K+ channel blockers (One way ANOVA), being fluoxetine and spadin TREK specific. Ct: control; Fl: fluoxetine; Sp: spadin

To ensure this current is driven by TREK channels, the isolated riluzole-activated current (300 µM) was recorded in the presence of specific channel blockers (Figure 5 C). We observed that the riluzole-induced current was insensitive to ZnCl_2_ (Figure 5 D), indicating that this current is not mediated by Ca^+2^ channels. On the contrary, the current was absent in the presence of BaCl_2_ (Figure 5 D), a broad-spectrum K^+^ channel blocker. Additionally, the riluzole-induced current was significantly reduced in the presence of both fluoxetine and spadin (Figure 5 D), two specific TREK channel blockers^19,50,51^, revealing the presence of TREK channels in mouse ICG neurons.

### Intracardiac ganglion neurons express the three members of the TREK subfamily

To further characterize the expression of TREK channels in ICG neurons, immunofluorescence was performed in tissue samples (Figure 6 A) to demonstrate the presence of the TREK subfamily in the ICG neurons. We found that the three members of this subfamily, TREK1, TREK2, and TRAAK are expressed in the ganglion in cells displaying big and round nuclei, typically considered to be neurons ^3^. Furthermore, immunofluorescence was also performed in primary ICG neuronal cultures (Figure 6 B) and the presence of the three members of the TREK subfamily was also found.

**Figure 6.**
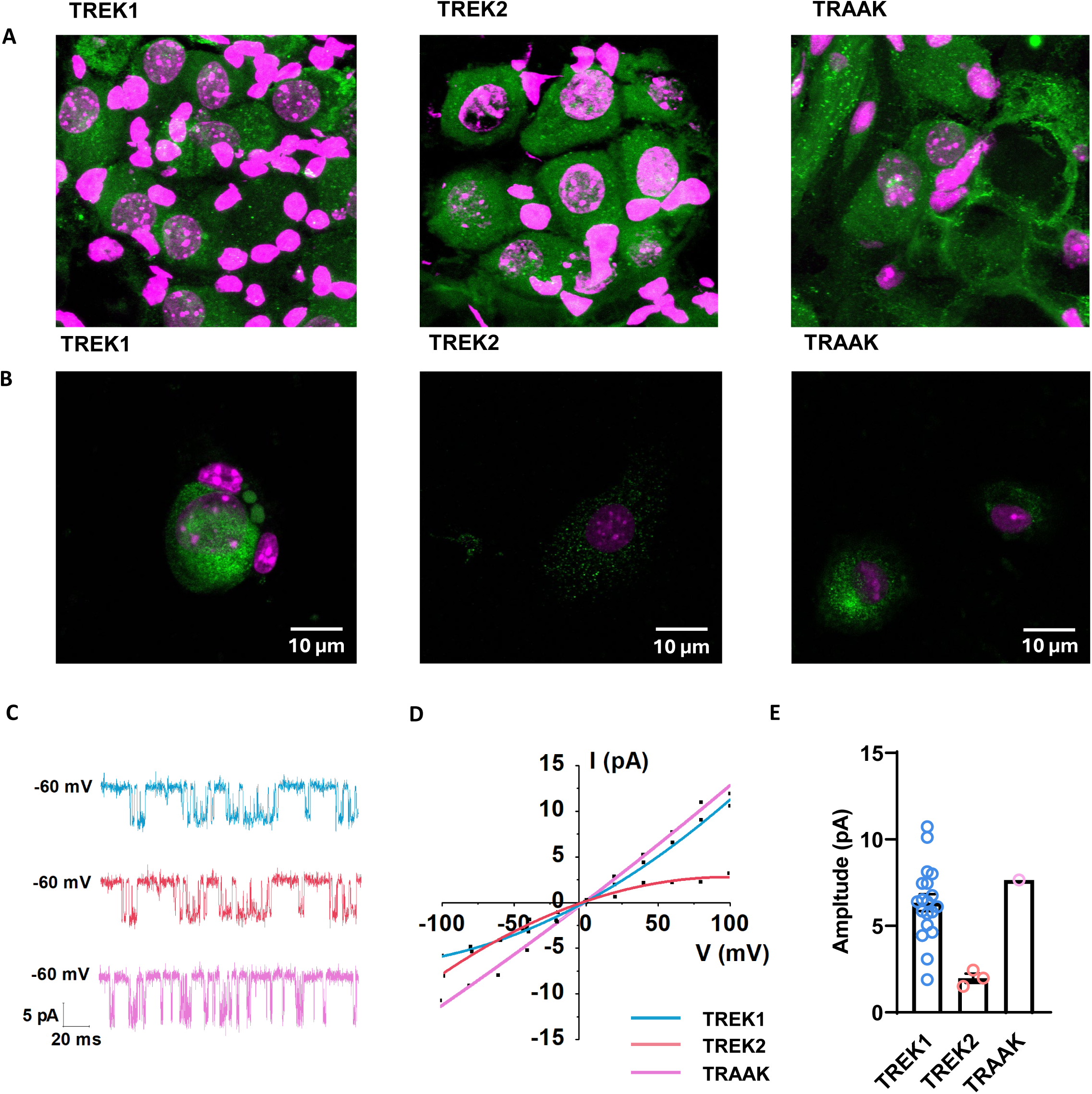
TREK channels are expressed in intracardiac ganglion neurons. (A) Immunofluorescence for TREK 1 (left), TREK2 (middle) and TRAAK (right) in tissue sections of the intracardiac ganglion (ICG). (B) Immunofluorescence for TREK 1 (left), TREK2 (middle) and TRAAK (right) in primary neuron cultures. Nuclei are shown in blue and the TREK subfamily shown in green. (C) Representative single channel recording of a TREK1 (red), TREK2 (blue) and TRAAK (green) channels and average voltage-current relationship and (D) amplitude of TREK1 (n=19), TREK 2 (n=3) and TRAAK (n=1), respectively.

Finally, the activity of individual TREK channels was assessed using cell-attached recordings under symmetric K^+^ conditions^52^, as observed in representative traces of the three subfamily members in Figure 6 C. Importantly, the single channels recorded in 19 of 23 cells presented a I-V with outward rectification (Figure 6 D, blue trace), typically associated to TREK1^53^, an amplitude of 6.34 ± 0.49 pA (Figure 6E) and a conductance of 66.06 ± 7.65 pS when clamped at −60 mV. 3 of 23 cells showed inward rectification (Figure 6 D, red trace), a typical TREK2 I-V^52^, an amplitude of 1.98 ± 0.27 pA (Figure 7 E) and a conductance of 70.71 ± 0.65 pS. Finally, 1 single cell showed an I-V associated to TRAAK^53^, with no rectification (Figure 6 D, green trace), an amplitude of −7.66 pA (Figure 7 E) and a conductance of 135.60 pS. These data indicate that the three TREK channel members can be detected in ICG neurons. However, most of the recorded channels were TREK1, suggesting that it might be expressed in higher amounts than the others.

Altogether, these data demonstrate the presence of TREK channels in the ICG neurons, suggesting that they might be playing an important homeostatic role in these neurons and therefore, in the regulation of the ICNS.

## DISCUSSION

Heart activity regulation is governed by the interplay between the sympathetic and parasympathetic branches of the autonomic ANS, operating under the command of higher centres. Within this intricate regulatory framework, the ICNS is a local circuit formed by a diffuse ganglionated plexus along the atrial surface, the ICG^54^. In this study, we have electrophysiologically characterized the ICG and investigated the presence of various ionic currents responsible for its well-functioning. Notably, we provide the first evidence that TREK channels contribute to RMP stabilization and excitability control of mouse ICG neurons.

The first step to start our neurochemical and electrophysiological study of the ICG neurons was to localize this nucleus in mouse tissue. The presence of these neuronal ganglia in murine hearts has been previously reported in the atrial region, atrioventricular (AV) groove, epicardium, interatrial septum and adipose tissue above the atria and around the aorta and veins^1,3,7,11,41,55^. Consistent with this, we found it in the AV groove of mouse hearts. These neurons were characterized by big, rounded somas perfectly visible with a Nissl staining. The ganglion size was quite heterogeneous in shape and neuronal content, ranging from 15 to 60 neurons. Although the presence of ICG has been also reported in ventricular regions^56^, we did not observe the presence of neurons in lower heart locations. The sinoatrial and AV nodes are innervated by this local circuit to control and regulate heart conduction^55^, thus, even though no somas were found in this region, it is possible that neuronal axons and prolongations innervate the lower regions of the heart.

The neurochemical profile of ICG neurons has been deeply studied^2,3,7,55^ and revealed a complex phenotype in mammalian cardiac neurons^2^. Although cholinergic expression is the main type reported in the ICG^2^, some neurons present tyrosine hydroxylase (TH)^2,3,7^ or adrenergic^33^ immunoreactivity. Calbindin is also expressed, normally co-stained with choline acetyltransferase (ChAT) or TH^2^, and a little neuronal population show immunoreactivity to nitric oxide synthases (NOS) and vesicular glutamate transporter 2 (VGLUT2). Neuropeptide Y (NPY) and cocaine and amphetamine-regulated transcript (CART) peptide, and nicotinic receptors have also been reported in these neurons^2,3^. We have reinsured the presence of ICG neurons thanks to the positive catecholaminergic phenotype (TH-positive neurons), and for the first time we have demonstrated that some of the ICG neurons are CR-positive, a typical characteristic of ganglionated neurons^37,57^.

From the electrophysiological point of view, we found that ICG neurons display a RMP of 70±1.2mV and a membrane capacitance of 33.9±1.4 pF, consistent with previous studies^2,3,11,27,41,58^. ICG neurons can be classified in three neuronal types based on their firing properties during a depolarizing step: phasic neurons, discharging 1 to 3 APs; adapting neurons, which adapt along the current pulse and discharge a maximum ∼ 6 APs; and tonic neurons, which discharge during the whole current pulse injection a maximum of ∼47 APs. Based on this classification, we found that adapting neurons are the predominant neuronal type (66%) followed by phasic (33%) and tonic (12%) neurons. Although our findings agree with previous studies ^8,47^ others have reported phasic neurons as the predominant neuronal type^2,3,11^ while tonic neurons are sometimes not even found^2,3^, probably due to insufficient sample sizes. Regarding AP individual parameters, the three neuronal types present similar passive and active properties, as previously shown^2^. However, we found differences in the frequency and threshold of APs, with tonic neurons exhibiting a significantly lower firing threshold compared to phasic and adapting neurons, likely contributing to higher firing frequency^3^.

Many cell types, including neurons in the ANS^38,39^ and cardiac nodal cells^59^ present TTX-resistant Na^+^ currents. However, our data shows that all the recorded cells show a clear sensitivity to TTX, demonstrating that this type of TTX-resistant currents are not present in mouse ICG neurons. Additionally, some studies point to a contribution of Ca^2+^ in the rise of AP in ICG rat neurons^43^, although this is controversial, since other reports suggest that it is uniquely mediated by Na^+ 3,41^. Our data show that AP generation is mediated by TTX-sensitive voltage-gated Na^+^ channels, supporting the idea that Ca^2+^ is not involved in the generation of AP in these cells. Furthermore, several K^+^ currents have a role in the intrinsic cardiac control of different species, including rat and guinea pig^8,12,42,60^. KCNQ and HCN channels, responsible for the I_M_ and I_H_, respectively, have been found in rat and guinea pig intrinsic cardiac neurons^8,61^. Additionally, Ca^2+^ voltage-dependent channels and K^+^ delayed rectifier channels, together with large conductance calcium-activated K^+^ channels (BK), which shape the AP in neonatal rat intracardiac neurons^10^, have been found in adult rats intracardiac neurons^62^. Our results show that mouse ICG neurons also express these channels, indicating that the activity of these neurons may be preserved among mammals.

TREK channels are critically implicated in heart physiology, being involved in the regulation of SA node excitability^22^ and cardiac AP duration^23^. However, their role on ICG neurons was unknown. We hypothesized that TREK channels may be present in the ICNS participating in the local regulation of the heart electrical conduction and motility. Importantly, most of the research studying the implication of TREK channels in heart function have been centred in TREK1^22,24,31,63,64^ probably due to its higher expression in many tissues and neuronal ganglion compared to the other members of the subfamily. By using both immunohistochemistry and electrophysiology, we demonstrate for the first time that the three members of the TREK subfamily (TREK1, TREK2 and TRAAK) are functionally expressed in ICG neurons in both heart tissue and ICG neuron primary cultures. We have shown that, in the presence of a cocktail of several blockers to isolate TREK currents, the addition of riluzole produced an outward current which was only inhibited by barium, a K^+^ channel blocker, and by the TREK channel blockers fluoxetine and spadin. In addition to that, the presence of TREK channels was strongly reinforced with the single channel experiments, where we observed the I-V curve of the three members, with a majority of TREK1 channels recorded.

In summary, the present results show that TREK channels are present in mouse ICG neurons and might be participating in RMP stabilization and excitability control. TREK channels have been associated to both heart physiology^22,31,64^ and pathophysiology with a possible neuroprotective role against ischemia, AF, arrhythmias, or cardiac hypertrophy^23–26,65^. Furthermore, the whole subfamily is modulated by temperature, changes in intracellular and extracellular pH or membrane stretc^66–71^, and it is possible all they are playing a neuroprotective role in the ICNS, as it has been demonstrated with TREK1^23,65^. Understanding how cardiac neurons work will shed light into the cellular mechanisms during physiological and pathological conditions, such as cardiac fibrillation and arrhythmia. Thus, our results unveil the presence of TREK channels and their possible role in neuroprotection.

## ACKNOWLEGEMENTS

The authors would like to thank the University of Vigo for the support and animal facilities technical assistance. We also thank the Galicia Sur Research Institute Microscopy Service for helping with the experiments. We thank all the members of the Neuro lab for useful discussions and for their invaluable support. This study was funded by the Spanish government (PID2022-138236NB-I00) to J.A.L.; Ministerio de Ciencia, Innovación y Universidades (MICIU), Agencia Estatal de Investigación (AEI) and European Social Fund plus ESF+ (RYC2022-035546-I) to A.C.; and by Programa de Axudas á etapa predoutoral da Consellería de Cultura, Educación e Universidades da Xunta de Galicia (ED481A-2022) to M.R-C.

## AUTHOR’S CONTRIBUTION

A.C-R., S.H-P. and M.R-C. performed electrophysiology. A.C-R., D.B-R. and A.R-T. performed immunohistochemistry. A.C-R., S. H-P. and A.C. analysed data. A.C-R., S.H-P., A.C. and J.A.L. conceptualized the study and wrote the manuscript. All authors read, edited and approved the last version of the manuscript.

## BIBLIOGRAPHY

1. Ashton, J. L. et al. Evidence of structural and functional plasticity occurring within the intracardiac nervous system of spontaneously hypertensive rats. Am J Physiol Heart Circ Physiol 318, (2020).

2. Lizot, G. et al. Molecular and functional characterization of the mouse intrinsic cardiac nervous system. Heart Rhythm 19, 1352–1362 (2022).

3. Hoard, J. L., Hoover, D. B. & Wondergem, R. Phenotypic properties of adult mouse intrinsic cardiac neurons maintained in culture. Am J Physiol Cell Physiol 293, 1875–1883 (2007).

4. Alves, F. H. F., Crestani, C. C. & Corrêa, F. M. A. The insular cortex modulates cardiovascular responses to acute restraint stress in rats. Brain Res 1333, 57–63 (2010).

5. Harper, R. M., Kumar, R., Macey, P. M., Ogren, J. A. & Richardson, H. L. Functional Neuroanatomy and Sleep-Disordered Breathing: Implications for Autonomic Regulation. Anat Rec 295, 1385–1395 (2012).

6. Oppenheimer, S. M., Gelb, A., Girvin, J. P. & Hachinski, V. C. Cardiovascular effects of human insular cortex stimulation. Neurology 42, 1727 (1992).

7. Rysevaite, K. et al. Morphologic pattern of the intrinsic ganglionated nerve plexus in mouse heart. Heart Rhythm 8, 448–454 (2011).

8. Cuevas, J., Harper, A. A., Trequattrini, C. & Adams, D. J. Passive and active membrane properties of isolated rat intracardiac neurons: Regulation by H- and M-currents. J Neurophysiol 78, 1890–1902 (1997).

9. Pérez, G. J., Desai, M., Anderson, S. & Scornik, F. S. Large-conductance calcium-activated potassium current modulates excitability in isolated canine intracardiac neurons. Am J Physiol Cell Physiol 304, (2013).

10. Franciolini, F. et al. Large-conductance calcium-activated potassium channels in neonatal rat intracardiac ganglion neurons. Pflügers Archiv 441, 629–638 (2001).

11. Rimmer, K. & Harper, A. A. Developmental Changes in Electrophysiological Properties and Synaptic Transmission in Rat Intracardiac Ganglion Neurons. J Neurophysiol 95, 3543–3552 (2006).

12. Allen, T. G. J. & Burnstock, G. The actions of adenosine 5′-triphosphate on guinea-pig intracardiac neurones in culture. Br J Pharmacol 100, 269–276 (1990).

13. Selga, E. et al. Molecular heterogeneity of large-conductance calcium-activated potassium channels in canine intracardiac ganglia. Channels 7, 322–328 (2013).

14. Cadaveira-Mosquera, A. et al. Expression of K2P Channels in Sensory and Motor Neurons of the Autonomic Nervous System. Journal of Molecular Neuroscience 48, 86–96 (2012).

15. Patel, A. & Honoré, E. The TREK two P domain K+ channels. Journal of Physiology 539, 647 (2002).

16. Ehling, P., Cerina, M., Budde, T., Meuth, S. G. & Bittner, S. The CNS under pathophysiologic attack—examining the role of K2P channels. Pflugers Arch 467, 959–972 (2015).

17. Djillani, A., Mazella, J., Heurteaux, C. & Borsotto, M. Role of TREK-1 in health and disease, focus on the central nervous system. Front Pharmacol 10, 1–15 (2019).

18. Spencer, K. A. et al. TREK-1 and TREK-2 Knockout Mice Are Not Resistant to Halothane or Isoflurane. Anesthesiology 139, 63–76 (2023).

19. Kennard, L. E. et al. Inhibition of the human two-pore domain potassium channel, TREK-1, by fluoxetine and its metabolite norfluoxetine. Br J Pharmacol 144, 821–829 (2005).

20. Duprat, F. et al. The neuroprotective agent riluzole activates the two P domain K+ channels TREK-1 and TRAAK. Mol Pharmacol 57, 906–912 (2000).

21. Cadaveira-Mosquera, A., Ribeiro, S. J., Reboreda, A., Pérez, M. & Lamas, J. A. Activation of TREK currents by the neuroprotective agent riluzole in mouse sympathetic neurons. Journal of Neuroscience 31, 1375–1385 (2011).

22. Unudurthi, S. D. et al. Two-Pore K+ channel TREK-1 regulates sinoatrial node membrane excitability. J Am Heart Assoc 5, (2016).

23. Kamatham, S., Waters, C. M., Schwingshackl, A. & Mancarella, S. TREK-1 protects the heart against ischemia-reperfusion-induced injury and from adverse remodeling after myocardial infarction. Pflugers Arch 471, 1263–1272 (2019).

24. Wang, W. et al. An Increased TREK-1–like Potassium Current in Ventricular Myocytes During Rat Cardiac Hypertrophy. J Cardiovasc Pharmacol 61, (2013).

25. Takahira, M., Sakurai, M., Sakurada, N. & Sugiyama, K. Fenamates and diltiazem modulate lipid-sensitive mechano-gated 2P domain K+ channels. Pflugers Arch 451, 474–478 (2005).

26. Herrera-Pérez, S., Rueda-Ruzafa, L., Campos-Ríos, A., Fernández-Fernández, D. & Lamas, J. A. Antiarrhythmic calcium channel blocker verapamil inhibits trek currents in sympathetic neurons. Front Pharmacol 13, (2022).

27. Hogg, R. C. et al. Mechanisms of verapamil inhibition of action potential firing in rat intracardiac ganglion neurons. Journal of Pharmacology and Experimental Therapeutics 289, 1502–1508 (1999).

28. Haipeng, Z., Shiyong, Y., Mackenzie, L. & Saint, D. A. Expression of 2-pore potassium channel Trek-1 in human heart. Eur. Heart J 26, (2005).

29. Fink, M. et al. Cloning, Functional Expression and Brain Localization of a Novel Unconventional Outward Rectifier K+ Channel. The EMBO Journal vol. 15 (1996).

30. Schmidt, C. et al. Cardiac expression and atrial fibrillation-associated remodeling of K2P2.1 (TREK-1) K+ channels in a porcine model. Life Sci 97, 107–115 (2014).

31. Wiedmann, F. et al. Mechanosensitive TREK-1 two-pore-domain potassium (K2P) channels in the cardiovascular system. Prog Biophys Mol Biol 159, 126–135 (2021).

32. Aimond, F., Rauzier, J. M., Bony, C. & Vassort, G. Simultaneous Activation of p38 MAPK and p42/44 MAPK by ATP Stimulates the K+ Current ITREK in Cardiomyocytes. Journal of Biological Chemistry 275, 39110–39116 (2000).

33. Horackova, M., Armour, J. A. & Byczko, Z. Distribution of intrinsic cardiac neurons in whole-mount guinea pig atria identified by multiple neurochemical coding. Cell Tissue Res 297, 409–421 (1999).

34. Foo, K. S., Hellysaz, A. & Broberger, C. Expression and colocalization patterns of calbindin-D28k, calretinin and parvalbumin in the rat hypothalamic arcuate nucleus. J Chem Neuroanat 61–62, 20–32 (2014).

35. Mandal, A. et al. A novel method for culturing enteric neurons generates neurospheres containing functional myenteric neuronal subtypes. J Neurosci Methods 407, 110144 (2024).

36. Masliukov, P. M., Moiseev, K., Budnik, A. F., Nozdrachev, A. D. & Timmermans, J.-P. Development of Calbindin- and Calretinin-Immunopositive Neurons in the Enteric Ganglia of Rats. Cell Mol Neurobiol 37, 1257–1267 (2017).

37. Barshack, I., Fridman, E., Goldberg, I., Chowers, Y. & Kopolovic, J. The loss of calretinin expression indicates aganglionosis in Hirschsprung’s disease. J Clin Pathol 57, 712–716 (2004).

38. Fernández-Fernández, D. et al. Activation of TREK currents by riluzole in three subgroups of cultured mouse nodose ganglion neurons. PLoS One 13, 1–24 (2018).

39. Campos, F. V., Moreira, T. H., Beirão, P. S. L. & Cruz, J. S. Veratridine modifies the TTX-resistant Na+ channels in rat vagal afferent neurons. Toxicon 43, 401–406 (2004).

40. Rugiero, F. et al. Selective Expression of a Persistent Tetrodotoxin-Resistant Na Current and Na1.9 Subunit in Myenteric Sensory Neurons. The Journal of Neuroscience 23, 2715 (2003).

41. Selyanko, A. A. Membrane properties and firing characteristics of rat cardiac neurones in vitro. J Auton Nerv Syst 39, 181–189 (1992).

42. Xi-Moy, S. X. & Dun, N. J. Potassium currents in adult rat intracardiac neurones. J Physiol 486, 15–31 (1995).

43. Xu, Z.-J. & Adams, D. J. Resting Membrane Potential and Potassium Currents in Cultures Parasympathetic Neurones from Rat Intracardiac Ganglia. Journal of Physiology vol. 456 (1992).

44. Mori, F., Tanji, K., Yoshimoto, M., Takahashi, H. & Wakabayashi, K. Demonstration of α-Synuclein Immunoreactivity in Neuronal and Glial Cytoplasm in Normal Human Brain Tissue Using Proteinase K and Formic Acid Pretreatment. Exp Neurol 176, 98–104 (2002).

45. Gu, X.-L. et al. Astrocytic expression of Parkinson’s disease-related A53T α-synuclein causes neurodegeneration in mice. Mol Brain 3, 12 (2010).

46. Tanji, K. et al. Expression of α-synuclein in a human glioma cell line and its up-regulation by interleukin-1β. Neuroreport 12, (2001).

47. Selyanko, A. A. et al. Two Types of K+ Channel Subunit, Erg1 and KCNQ2/3, Contribute to the M-Like Current in a Mammalian Neuronal Cell. The Journal of Neuroscience 19, 7742 (1999).

48. Wang, H.-S. et al. KCNQ2 and KCNQ3 Potassium Channel Subunits: Molecular Correlates of the M-Channel. Science (1979) 282, 1890–1893 (1998).

49. Fang, T. et al. Stage at which riluzole treatment prolongs survival in patients with amyotrophic lateral sclerosis: a retrospective analysis of data from a dose-ranging study. Lancet Neurol 17, 416–422 (2018).

50. Kobayashi, T., Washiyama, K. & Ikeda, K. Inhibition of G protein-activated inwardly rectifying K+ channels by fluoxetine (Prozac). Br J Pharmacol 138, 1119–1128 (2003).

51. Mazella, J. et al. Spadin, a Sortilin-Derived Peptide, Targeting Rodent TREK-1 Channels: A New Concept in the Antidepressant Drug Design. PLoS Biol 8, e1000355-(2010).

52. Han, J., Truell, J., Gnatenco, C. & Kim, D. Characterization of four types of background potassium channels in rat cerebellar granule neurons. J Physiol 542, 431–444 (2002).

53. Blin, S. et al. Mixing and matching TREK/TRAAK subunits generate heterodimeric K2P channels with unique properties. Proceedings of the National Academy of Sciences 113, 4200–4205 (2016).

54. Durães Campos, I., Pinto, V., Sousa, N. & Pereira, V. H. A brain within the heart: A review on the intracardiac nervous system. J Mol Cell Cardiol 119, 1–9 (2018).

55. Pauza, D. H. et al. Neuroanatomy of the murine cardiac conduction system. A combined stereomicroscopic and fluorescence immunohistochemical study. Auton Neurosci 176, 32–47 (2013).

56. Armour, J. A., Murphy, D. A., Yuan, B. X., Macdonald, S. & Hopkins, D. A. Gross and microscopic anatomy of the human intrinsic cardiac nervous system. Anatomical Record 247, 289–298 (1997).

57. Musser, M. A., Correa, H. & Southard-Smith, E. M. Enteric Neuron Imbalance and Proximal Dysmotility in Ganglionated Intestine of the Sox10Dom/+ Hirschsprung Mouse Model. Cell Mol Gastroenterol Hepatol 1, 87–101 (2015).

58. Catacuzzeno, L. & Adams, D. J. Mechanisms of Verapamil Inhibition of Action Potential Firing in Rat Intracardiac Ganglion Neurons. Article in Journal of Pharmacology and Experimental Therapeutics http://www.jpet.org (1999).

59. Lei, M. et al. Requirement of neuronal- and cardiac-type sodium channels for murine sinoatrial node pacemaking. J Physiol 559, 835–848 (2004).

60. Hassall, C. J. S. & Burnstock, G. Intrinsic neurones and associated cells of the guinea-pig heart in culture. Brain Res 364, 102–113 (1986).

61. Arichi, S., Sasaki-Hamada, S., Kadoya, Y., Ogata, M. & Ishibashi, H. Excitatory effect of bradykinin on intrinsic neurons of the rat heart. Neuropeptides 75, 65–74 (2019).

62. Whyte, K. A., Hogg, R. C., Dyavanapalli, J., Harper, A. A. & Adams, D. J. Reactive oxygen species modulate neuronal excitability in rat intrinsic cardiac ganglia. Auton Neurosci 150, 45–52 (2009).

63. Unudurthi, S. D. et al. Two-Pore K+ channel TREK-1 regulates sinoatrial node membrane excitability. J Am Heart Assoc 5, (2016).

64. Li, X. T. et al. The stretch-activated potassium channel TREK-1 in rat cardiac ventricular muscle. Cardiovasc Res 69, 86–97 (2006).

65. Buckler, K. J. & Honoré, E. The lipid-activated two-pore domain K+ channel TREK-1 is resistant to hypoxia: implication for ischaemic neuroprotection. J Physiol 562, 213–222 (2005).

66. Kang, D., Choe, C. & Kim, D. Thermosensitivity of the two-pore domain K+ channels TREK-2 and TRAAK. J Physiol 564, 103–116 (2005).

67. Maingret, F. TREK-1 is a heat-activated background K+ channel.Maingret, F. (2000). TREK-1 is a heat-activated background K+ channel. The EMBO Journal, 19(11), 2483–2491. 10.1093/emboj/19.11.2483The EMBO Journal 19, 2483–2491 (2000).

68. Honoré, E., Maingret, F., Lazdunski, M. & Patel, A. J. An intracellular proton sensor commands lipid- and mechano-gating of the K+ channel TREK-1. EMBO J 21, 2968-2976–2976 (2002).

69. Maingret, F., Patel, A. J., Lesage, F., Lazdunski, M. & Honoré, E. Mechano- or acid stimulation, two interactive modes of activation of the TREK-1 potassium channel. Journal of Biological Chemistry 274, 26691–26696 (1999).

70. Zhang, H., Shepherd, N. & Creazzo, T. L. Temperature-sensitive TREK currents contribute to setting the resting membrane potential in embryonic atrial myocytes. J Physiol 586, 3645–3656 (2008).

71. Reyes, R. et al. Cloning and Expression of a Novel pH-sensitive Two Pore Domain K+ Channel from Human Kidney. Journal of Biological Chemistry 273, 30863–30869 (1998).

